# Regulation of spontaneous neurotransmission and homeostatic synaptic plasticity by synaptotagmin-1 disease variants at the SNARE primary interface

**DOI:** 10.64898/2026.02.17.706274

**Authors:** Elena D. Bagatelas, Ok-ho Shin, Reagan M. Armstrong, Qiangjun Zhou, Ege T. Kavalali

**Affiliations:** Department of Pharmacology, Vanderbilt University, Nashville, TN 37240-7933, USA; Vanderbilt Brain Institute, Vanderbilt University, Nashville, TN 37240-7933, USA; Department of Cell and Developmental Biology, Vanderbilt University, Nashville, TN 7240-7933, USA; Vanderbilt Center for Structural Biology, Vanderbilt University, Nashville, TN 37240-7933, USA

## Abstract

*De novo* mutations in synaptotagmin-1 (syt1) cause a rare neurodevelopmental disorder, manifesting in global developmental delay, ophthalmic abnormalities, infantile hypotonia, facial dysmorphisms, absent speech, EEG abnormalities, and hyperkinetic movements, ranging from moderate to severe. Here, we evaluate eleven patient-relevant mutations spanning the Ca^2+^ binding domains of syt1—C2A and -C2B impact neurotransmission. We found that the mutation causing the most severe impact on neurotransmission, p.N341S, triggers aberrant spontaneous neurotransmission and occludes homeostatic synaptic plasticity signaling pathways. Our results suggest that potential phosphorylation of this newly introduced Ser residue underlies the functional change. A serine missense mutation creates a novel phosphorylation site as a broad spectrum protein kinase inhibitor rescues spontaneous neurotransmission. We identify key residues, localized to the primary interface between syt1 and SNAP-25, responsible for this shift in syt1 function in synaptic vesicle release. Substituting neutral amino acids at residue 341 alters the interaction of the Ser mutation, with double mutations in the surrounding amino acids in the primary interface rescuing synaptic function. These results provide a framework for how a syt1 point mutation introduces a substrate for phosphorylation and disrupts intermolecular interactions at the primary interface with SNAP-25 altering spontaneous neurotransmission and homeostatic plasticity.

**AUTHOR SUMMARY:** Mutations in synaptotagmin-1 (*SYT1*), a protein essential for communication between neurons, cause a rare neurodevelopmental disorder marked by developmental delay, low muscle tone, abnormal movements, vision problems, and disrupted brain electrical activity. Disease severity varies, and how specific syt1 mutations alter brain signaling remains unclear. In this study, we examined 11 disease-associated syt1 mutations that affect regions of the protein responsible for sensing calcium, a key trigger for neurotransmitter release. Using a range of electrophysiological approaches, we measured how these mutations influence different modes of synaptic communication within neuronal networks. We found that one mutation, N341S, produced the most severe disruption. Neurons carrying this mutation released neurotransmitters abnormally at rest and were unable to engage normal homeostatic plasticity mechanisms that stabilize brain activity. These effects suggest a fundamental breakdown in how synapses regulate signaling strength.

We investigated the molecular basis of this dysfunction and identified a likely explanation: the N341S mutation introduces a new serine residue that can be phosphorylated, a common regulatory modification in cells. Our data indicate that this newly created phosphorylation site alters syt1 function, as blocking phosphorylation pathways could modify the mutant’s effects. Importantly, we also show that N341 residue lies within a critical interaction interface between syt1 and another synaptic protein, SNAP-25. Adjusting nearby amino acids to neutralize this interaction restores wild-type levels of synaptic signaling.

Together, these findings reveal how a single disease-associated mutation can rewire synaptic regulation by introducing a phosphorylation site, offering new insight into the underpinning of syt1-related neurodevelopmental disorders and potential therapeutic targets.

## INTRODUCTION

Neurotransmission underlies signaling across local and global networks in the brain. Dysfunction of this process can impair synapse physiology and network plasticity, with the canonical SNARE complex serving as essential machinery for neurotransmitter release^1–4^. In the last decade, advances made in genomics has led to the discovery of *de novo* mutations associated with SNARE-regulatory proteins that manifest neurodevelopmental disorders^5–7^. Much of our understanding of the etiology of these disorders comes from studies of patient-derived mutations, which reveal how indispensable vesicular machinery is for synaptic function^5^. Notably, the SNARE complex and its associated proteins are highly evolutionarily conserved. Therefore, elucidating how *de novo* mutations in this tightly coupled machinery give rise to a broad and multifaceted spectrum of clinical symptoms will provide important insight into disease mechanisms and inform the development of future therapeutic strategies.

Calcium (Ca^2+^) regulated exocytosis of vesicles depends on the syt1 protein, which acts in combination with the SNARE complex for efficient neurotransmitter release^8–10^. There are many syt protein isoforms expressed within the central nervous system that have diverse processes (syt1-syt17), with syt1 being the predominant isoform expressed in the mammalian cortex at both excitatory and inhibitory synapses. Syt1 is involved in both spontaneous and evoked neurotransmitter release, both critical in synaptic signaling^11–13^. Syt1 triggers vesicle exocytosis within milliseconds upon its C2A and C2B domains binding Ca^2+^ ions that enter the presynaptic terminal with an action potential stimulus^14,15^. Tethering of syt1 to the plasma membrane of vesicles occurs through its N-terminal transmembrane domain, while Ca^2+^ binding is mediated by the C2A and C2B domains at the protein’s C-terminal end^16–22^.

Baker Gordon Syndrome (BAGOS), or Synaptotagmin-1 (syt1) associated neurodevelopmental disorder, is considered an ultra-rare disorder with a prevalence of 1:1,000,000^23,24^. It is derived from mutations in the *SYT1* gene, which encodes the SNARE-associated regulatory protein synaptotagmin-1. BAGOS is associated with global developmental delay, ophthalmic abnormalities, infantile hypotonia, facial dysmorphisms, absent speech, EEG abnormalities, and hyperkinetic movements^6,25–27^. To date nearly 40 patients have been diagnosed with BAGOS since its initial classification in 2015, with only a small number of studies in the literature, three of which are individual case reports^19,20^. Individuals with this diagnosis harbor *de novo* mutations. There is currently no targeted treatment for BAGOS, with current efforts focusing on symptom management. Despite the seminal work on clinical phenotyping and basic functional studies on several variants, the underlying mechanistic correlations that manifest in this neurodevelopmental disorder have not been elucidated. Because the disease relevance of altered release patterns in this model is unclear, we focus on spontaneous neurotransmission, which is essential for synapse maturation and plasticity in particular during development. We have focused on how variants in the calcium-sensing region of the protein synaptotagmin-1 lead to a genetically determined neurodevelopmental disorder with high penetrance (Table 1). To mimic the human syt1 mutations that result in neurodevelopmental disorder, we recapitulated disease-relevant neuronal networks in rat hippocampal cultures to analyze functional consequences of syt1 mutations on synaptic transmission, network development and plasticity, and to test pharmacological approaches for rescuing elements of neurotransmission.

**Table 1:**
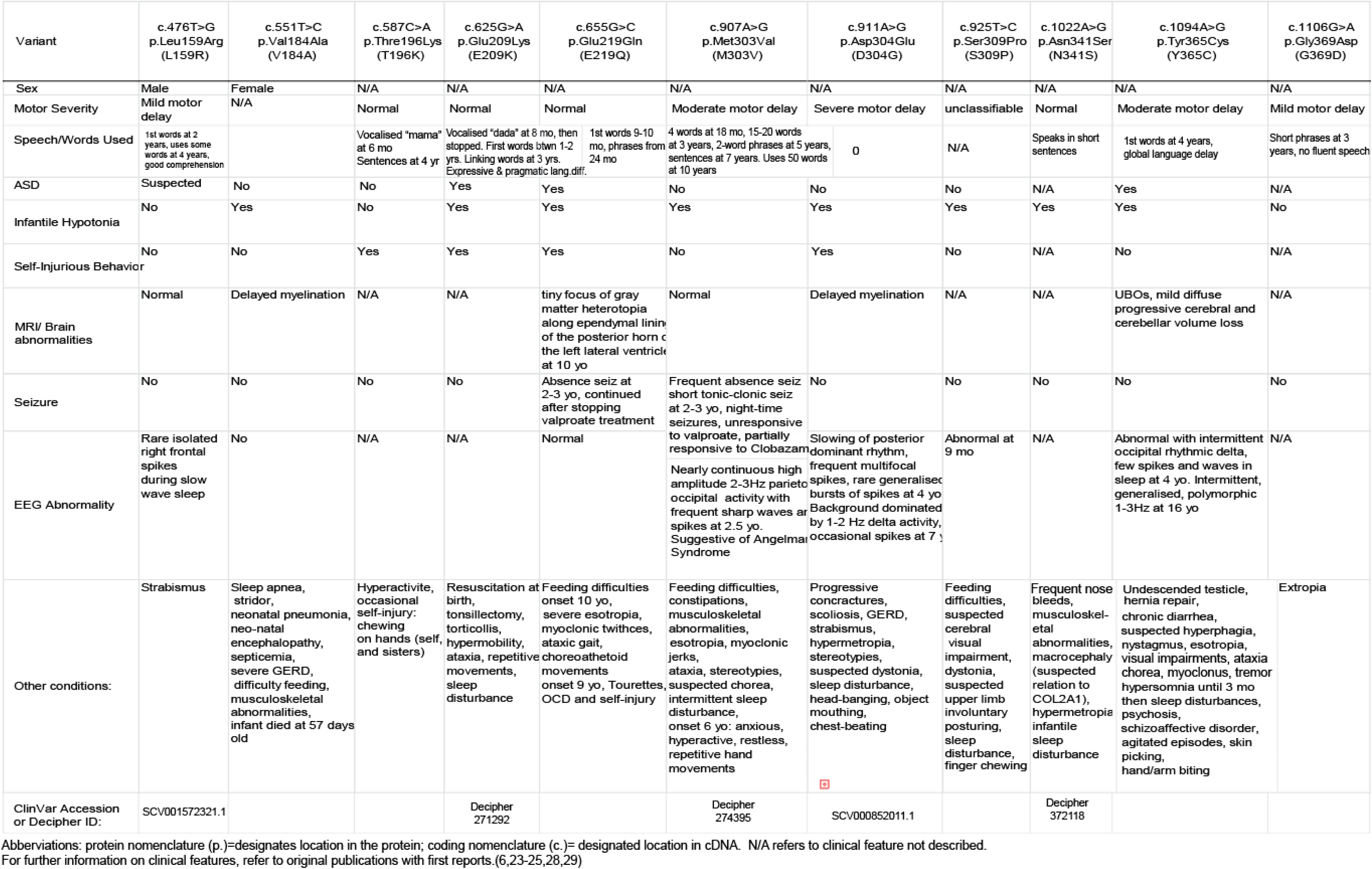
Clinical features of Syt1-associated neurodevelopmental disorder patient variants in this study.

## RESULTS

### Expression of select C2B variants on WT background augments spontaneous neurotransmission

Of the 11 syt1 variants, 5 are located along the C2A domain and 6 are located along the C2B domain (Fig. 1A) ^6,24,28,29^. All of the patient-relevant mutations we studied are *de novo* heterozygous missense mutations. We first looked at whether spontaneous synaptic transmission is altered by their expression. We created neuronal networks with the variant syt1 expressed on a wild-type (WT) background (Fig. 1B). Compared to WT syt1 overexpression, expression of all 5 of the C2A human variants did not alter spontaneous post-synaptic currents, including miniature excitatory (mEPSCs) or miniature inhibitory (mIPSCs) transmission (Fig. 1 C-H). However, 3 of the 6 C2B variants—D304G, S309P, and N341S—significantly augmented mEPSC transmission, with S309P and D304G increasing mEPSC amplitude, and N341S increasing both amplitude and frequency (Fig. 1 I-K). Notably, we also observed the N341S variant significantly affected inhibitory transmission, demonstrating augmented mIPSC amplitude and frequency (Fig. 1 M-O).

**Figure 1:**
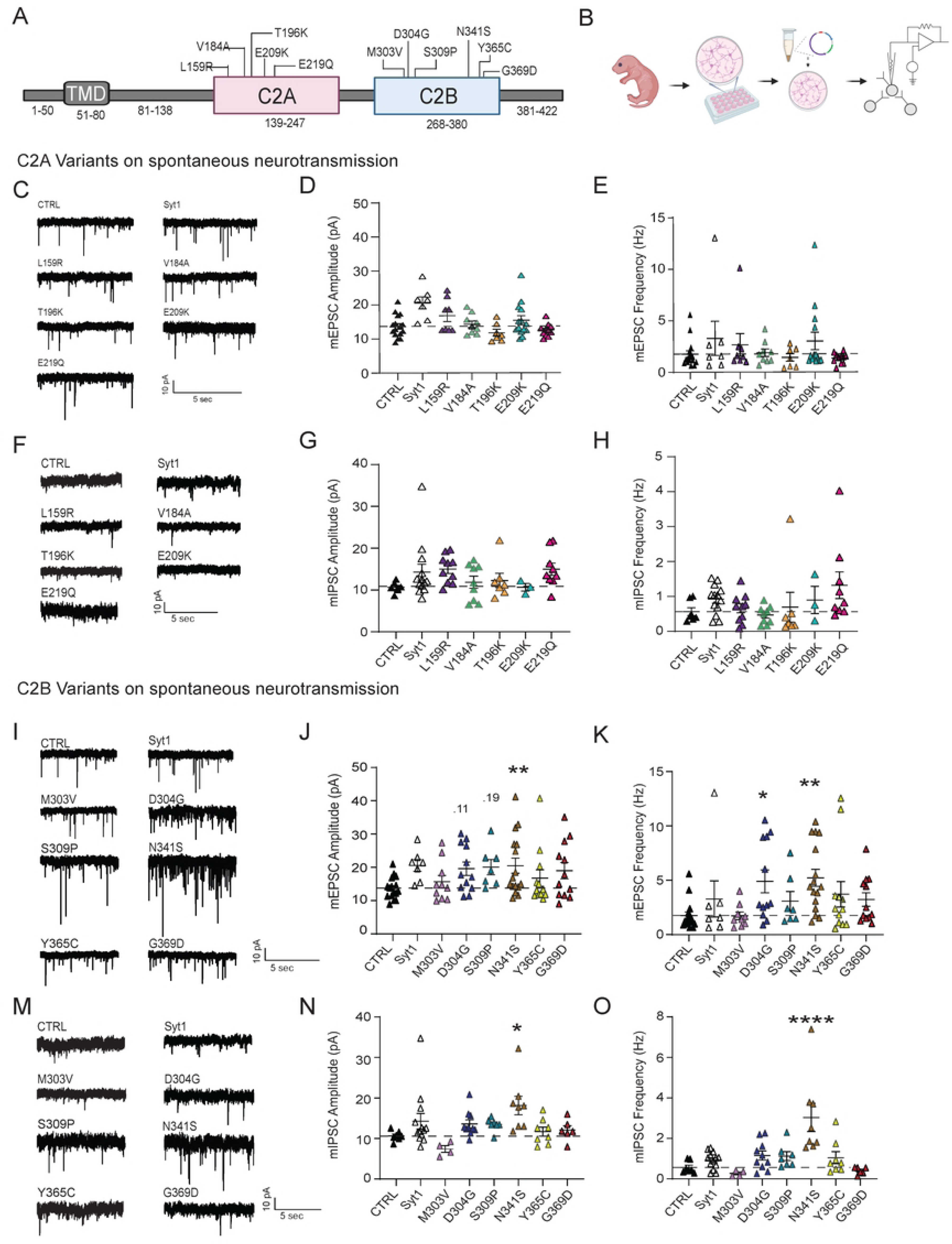
Effects of Syt1 variants on excitatory and inhibitory synaptic transmission. (A) Schematic of Syt1 protein segment with designated variants along the C2A and C2B domains. (N.B.: in these and all subsequent experiments, amino acid (aa) numbers pertain to the human syt1 sequences). (B) Experimental design: lentiviral constructs were transfected on DIV 4 with virus carrying empty vector (CTRL) as the positive control, Syt1 overexpression (Syt1), or Syt1 variants. Experiments were performed between DIV 14-21. (C) Representative raw traces of baseline mEPSCs on CTRL, Syt1 overexpression (Syt1), and Syt1 variants in the C2A domain. (D) Comparison of baseline mEPSC amplitudes (pA) (One-way ANOVA with Dunnett’s multiple comparisons test: CTRL vs. Syt1 overexpression p=0.1290; CTRL vs. L159R p=0.922; CTRL vs. V184A p>0.999; CTRL vs. T196K p=0.998; CTRL vs. E209K p=0.997; CTRL vs. E219Q p>0.999, N=2-4, n=7-19 per group). (E) Comparison of mEPSC frequencies (Hz) (One-way ANOVA with Dunnett’s multiple comparisons test: CTRL vs. Syt1 overexpression p=0.866; CTRL vs. L159R p=0.996; CTRL vs. V184A p>0.999; CTRL vs. T196K p>0.999; CTRL vs. E209K p=0.857; CTRL vs. E219Q p>0.999; N=2-4, n=7-19 per group). (F) Representative raw traces of baseline mEPSCs on CTRL and Syt1 variants in the C2B domain. (G) Comparison of baseline mEPSC amplitudes (pA) (One-way ANOVA with Dunnett’s multiple comparisons test: CTRL vs. M303V p=0.996; CTRL vs. D304G p=0.117; CTRL vs. S309P p=0.196; CTRL vs. N341S p=0.021; CTRL vs. Y365C p=0.856; CTRL vs. G369D p=0.210, N=2-5, n=7-16 per group). (H) Comparison of mEPSC frequencies (Hz) (One-way ANOVA with Dunnett’s multiple comparisons test: CTRL vs. M303V p>0.999; CTRL vs. D304G p=0.021; CTRL vs. S309P p=0.944; CTRL vs. N341S p=0.002; CTRL vs. Y365C p=0.382; CTRL vs. G369D p=0.743, N=2-5, n=7-16 per group). (I) Representative raw traces of baseline mIPSCs on CTRL, syt1 overexpression (Syt1), and Syt1 variants in the C2A domain. (J) Comparison of baseline mIPSC amplitudes (pA) (One-way ANOVA with Dunnett’s multiple comparisons test: CTRL vs. Syt1 overexpression p=0.352; CTRL vs. L159R p=0.276; CTRL vs. V184A p=0.999; CTRL vs. T196K p=0.996; CTRL vs. E209K p>0.999; CTRL vs. E219Q p=0.306, N=1-4, n=3-13 per group). (K) Comparison of mIPSC frequencies (Hz) (One-way ANOVA with Dunnett’s multiple comparisons test: CTRL vs. Syt1 overexpression p=0.977; CTRL vs. L159R p>0.999; CTRL vs. V184A p>0.999; CTRL vs. T196K p>0.999; CTRL vs. E209K p=0.999; CTRL vs. E219Q p=0.448, N=1-4, n=3-13 per group). (L) Representative raw traces of baseline mIPSCs in CTRL and select Syt1 variants in the C2B domain. (M) Comparison of baseline mIPSC amplitudes (pA) (One-way ANOVA with Dunnett’s multiple comparisons test: CTRL vs. M303V p=0.863; CTRL vs. D304G p=0.725; CTRL vs. S309P p=0.8462; CTRL vs. N341S p=0.010; CTRL vs. Y365C p>0.999; CTRL vs. G369D p=0.999, N=1-5, n=4-13 per group). (N) Comparison of mIPSC frequencies (Hz) for each group (One-way ANOVA with Dunnett’s multiple comparisons test: CTRL vs. M303V p=0.999; CTRL vs. D304G p=0.723; CTRL vs. S309P p=0.844; CTRL vs. N341S p<0.0001; CTRL vs. Y365C p=0.915; CTRL vs. G369D p>0.999, N=1-5, n=4-13 per group).

### Expressing syt1 variants on WT background has no significant effect on evoked neurotransmission

We expressed all 11 syt1 variants on a WT background and performed experiments using a bilateral stimulation probe to measure the effects on evoked inhibitory post-synaptic currents (eIPSC), such as on synaptic strength and presynaptic release probability (Fig 2A, B, E.) There were no changes observed in peak amplitude of eIPSCs with the 10 Hz stimulation protocol across all C2A or C2B variants (Fig. 2C, F.) There was also no change in paired pulse ratios (a measure of release probability), which corresponds to the response of the 2^nd^ stimulation divided by the 1^st^ (Fig. 2D, G.) Experiments were also performed with 20 Hz stimulation protocols (Supp. 1).

**Figure 2:**
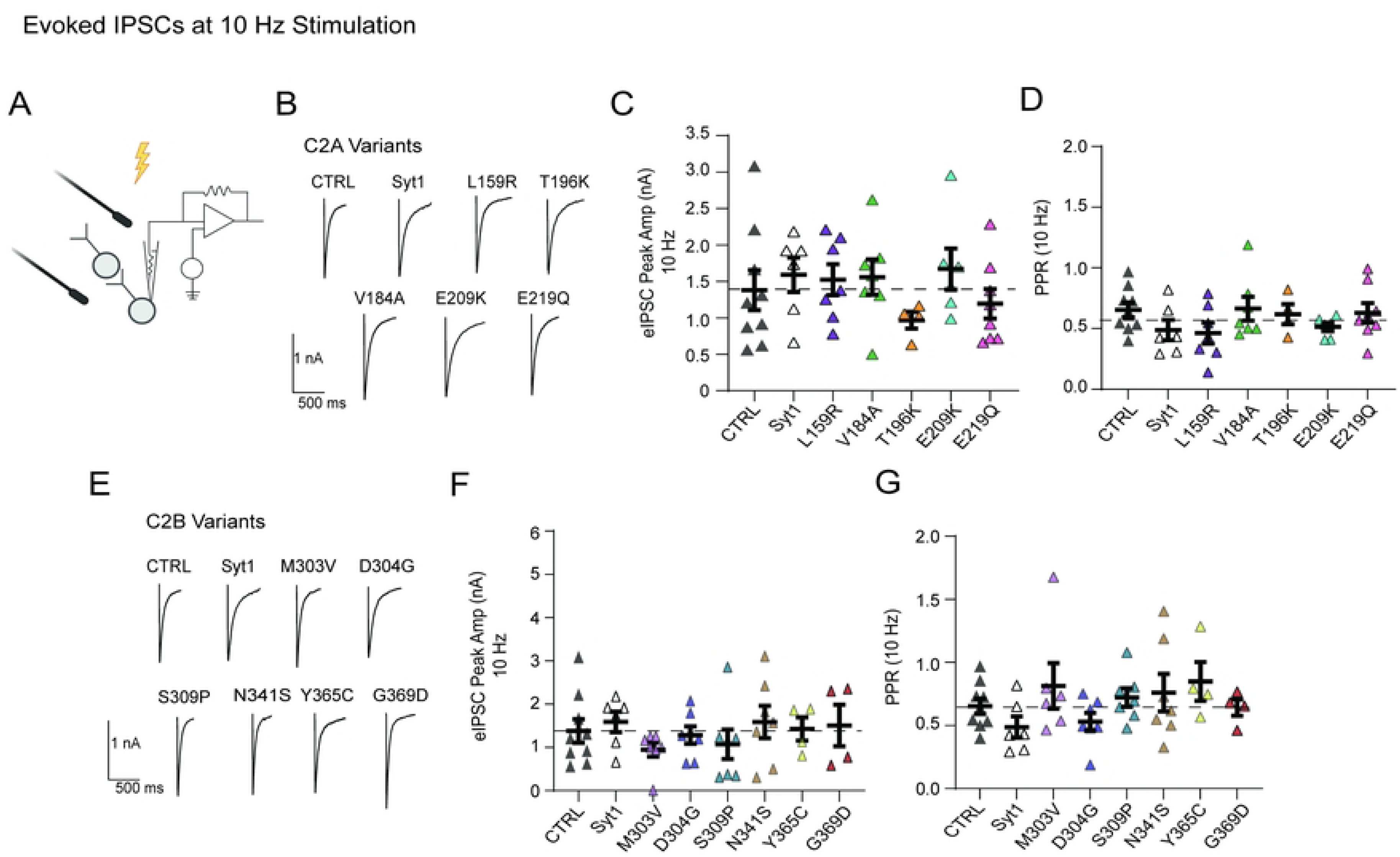
Expressing syt1 variants has no significant effect on evoked inhibitory neurotransmission. (A) Schematic illustrating stimulation of neuronal network to record evoked neurotransmission. (B) Representative evoked inhibitory current traces of empty vector (CTRL) and expressing C2A syt1 variants. (C) Comparison of the mean eIPSC amplitude (nA) at 10 Hz stimulation across C2A variants (Two-way ANOVA with Dunnett’s multiple comparisons test: CTRL vs. Syt1 OE p=0.999; CTRL vs. L159R p>999; CTRL vs. V184A p>0.999; CTRL vs. T196K p=0.967; CTRL vs. E209K p=0.993; CTRL vs. E219Q p=0.999; N=1-3, n=4-8 per group). (D) Comparison of paired-pulse ratio (PPR) at 10 Hz stimulation for each group (One-way ANOVA with Dunnett’s multiple comparisons test: CTRL vs. Syt1 OE p=0.967; CTRL vs. L159R p=.695; CTRL vs. V184A p>0.999; CTRL vs. T196K p>0.999; CTRL vs. E209K p=0.949; CTRL vs. E219Q p>0.999; N=1-3, n=4-8 per group). (E) Representative evoked inhibitory current traces of empty vector (CTRL) and expressing of C2B syt1 variants. (F) Comparison of mean eIPSC amplitude (nA) at 10 Hz stimulation across C2B variants (Two-way ANOVA with Dunnett’s multiple comparisons test: CTRL vs. Syt1 OE p=0.999; CTRL vs. M303V p=.877; CTRL vs. D304G p>0.999; CTRL vs. S309P p=0.986; CTRL vs. N341S p=0.999; CTRL vs. Y365C p>0.999; CTRL vs. G369D p>0.999; N=1-3, n=4-8 per group). (G) Comparison of paired-pulse ratio (PPR) at 10 Hz stimulation for each group (One-way ANOVA with Dunnett’s multiple comparisons test: CTRL vs. Syt1 OE p=0.867; CTRL vs. M303V p=0.881; CTRL vs. D304G p=0.972; CTRL vs. S309P p=0.999; CTRL vs. N341S p=0.990; CTRL vs. Y365C p>0.843; CTRL vs. G369D p>0.999; N=1-3, n=4-8 per group).

### N341S mutation drives further loss-of-function on syt1KD background

We did not observe any robust electrophysiological signatures with the C2A variants in our expression experiments on WT background. However, three of the C2B variants enhanced spontaneous excitatory transmission to varying degrees. To assess how these three variants affect synaptic mechanisms, we examined their phenotypes on a syt1 loss-of-function (LOF) background. Endogenous syt1 was knocked-down (KD) using shRNA lentivirus, and function was rescued using lentiviral expression of human D304G, S309P or N341S variants (Fig. 3A). Corroborating previous findings ^30^, we observed syt1KD significantly increased spontaneous neurotransmission through augmented mEPSC and mIPSC frequency (Fig. 3B, E)(27). We also observed all three C2B variants— D304G, S309P and N341S—increased both mEPSC and mIPSC amplitude (Fig. 3C, F). Notably, expression of the N341S variant on a syt1KD background further increased mEPSC and mIPSC frequency beyond that of syt1KD alone, suggesting that this mutation severely dysregulates spontaneous excitatory neurotransmission (Fig. 3D, G). Other syt1 variants were also tested on syt1 LOF background. Both mEPSC amplitude and frequency returned to WT levels after rescuing with all the C2A variants; C2B variants M303V, Y365C, and G369D returned to WT levels as well (Supp. 2). All C2A variants maintained mIPSC amplitudes similar to that of WT, except T196K which increased the amplitude. For mIPSC frequency, the C2A variants V184A and T196K sustained increased levels with all other C2A variants rescuing to WT levels. The C2B variant Y365C also increased mIPSC amplitude and frequency, whereas all the other C2B variants maintained WT levels for amplitude and frequency (Supp. 3).

**Figure 3:**
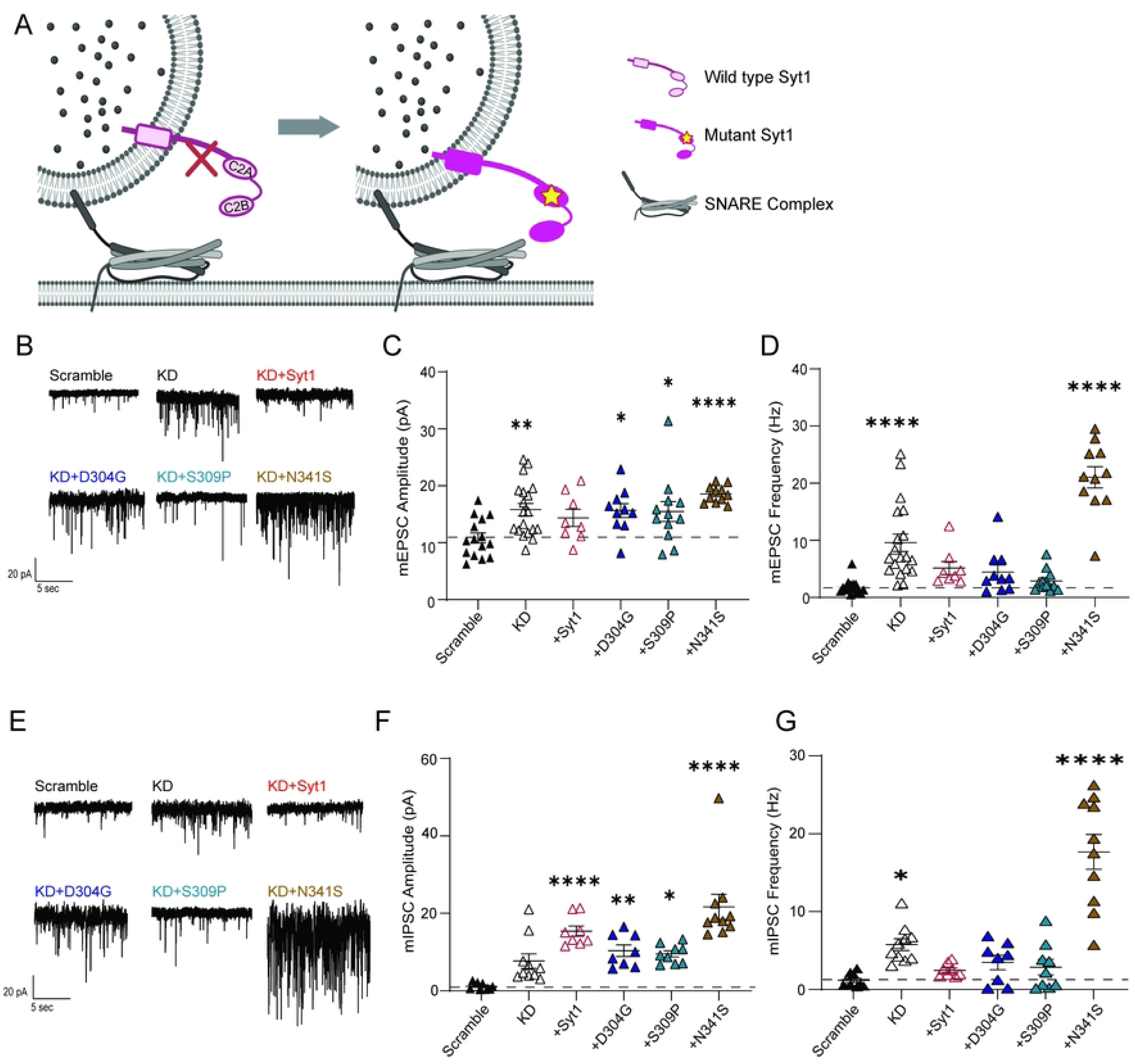
N341S mutation drives further loss-of-function on syt1KD background. (A) Schematic illustrating knockdown of syt1 and rescuing with syt1 variants. (B) Representative raw traces of baseline mEPSCs for scramble (negative control), Syt1 knockdown (KD), Syt1 rescue (Syt1), and select Syt1 variants in the C2B domain. (C) Comparison of baseline mEPSC amplitudes (pA) (One-way ANOVA with Dunnett’s multiple comparisons test: SCR vs KD p=0.004; SCR vs Syt1 rescue p=0.350; SCR vs. D304G p=0.039; SCR vs. S309P p=0.032; SCR vs. N341S p<0.0001, N=2-5, n=8-19 per group). (D) Comparison of baseline mEPSC frequency (Hz) (One-way ANOVA with Dunnett’s multiple comparisons test: SCR vs KD p<0.0001; SCR vs Syt1 rescue p=0.406; SCR vs. D304G p=0.626; SCR vs. S309P p=0.998; SCR vs. N341S p<0.0001, N=2-5, n=8-19 per group). (E) Representative raw traces of baseline mIPSCs for scramble (SCR), Syt1 knockdown (KD), Syt1 rescue (Syt1), and select Syt1 variants in the C2B domain. (F) Comparison of baseline mIPSC amplitudes (pA) (One-way ANOVA with Dunnett’s multiple comparisons test: SCR vs KD p=.054; SCR vs Syt1 rescue p>.0001; SCR vs. D304G p=0.005; SCR vs. S309P p=0.010; SCR vs. N341S p<0.0001, N=2-5, n=8-10 per group). (G) Comparison of baseline mIPSC frequencies (Hz) (One-way ANOVA with Dunnett’s multiple comparisons test: SCR vs KD p=0.027; SCR vs Syt1 rescue p=0.915; SCR vs. D304G p=0.549; SCR vs. S309P p=0.766; SCR vs. N341S p<0.0001, N=2-5, n=8-10 per group).

### Cell autonomous expression of syt1 variants reveal presynaptic changes elicit chronic changes in spontaneous neurotransmission

Thus far, we observed the most robust effects on spontaneous neurotransmission across the three C2B human variants—D304G, S309P, and N341S—with N341S increasing both amplitude and frequency, regardless of whether expressed on WT- or syt1KD-background. We next sought to determine whether the sustained increase in amplitude results from local presynaptic changes or from chronic network-wide alterations affecting both pre- and post-synaptic sites. To address this, we sparsely transfected neurons (∼20% of culture) with constructs co-expressing mEGFP in neurons expressing wild-type (syt1), D304G, S309P or N341S (Fig. 4A). This design consists of neurons with both wild-type (unlabeled) and syt1-variant-expressing neurons (GFP labeled) (Fig. 4B). Recordings from sparsely transfected cells expressing each of the 3 disease variants mirrored wild-type networks, with no change in mEPSC amplitude and frequency (Fig. 4C, D). This result suggests that the sustained increase in spontaneous mEPSC amplitude originates from presynaptic changes, where syt1-variant interferes with wild-type syt1 function when both are abundantly present at the same synapse. Alongside our syt1KD neurons rescued with the variants, these results support the earlier proposal that the amount of variant protein influences the spontaneous transmission phenotype, with increased levels of variant syt1, especially N341S, leading to deficits in spontaneous release^5^.

**Figure 4:**
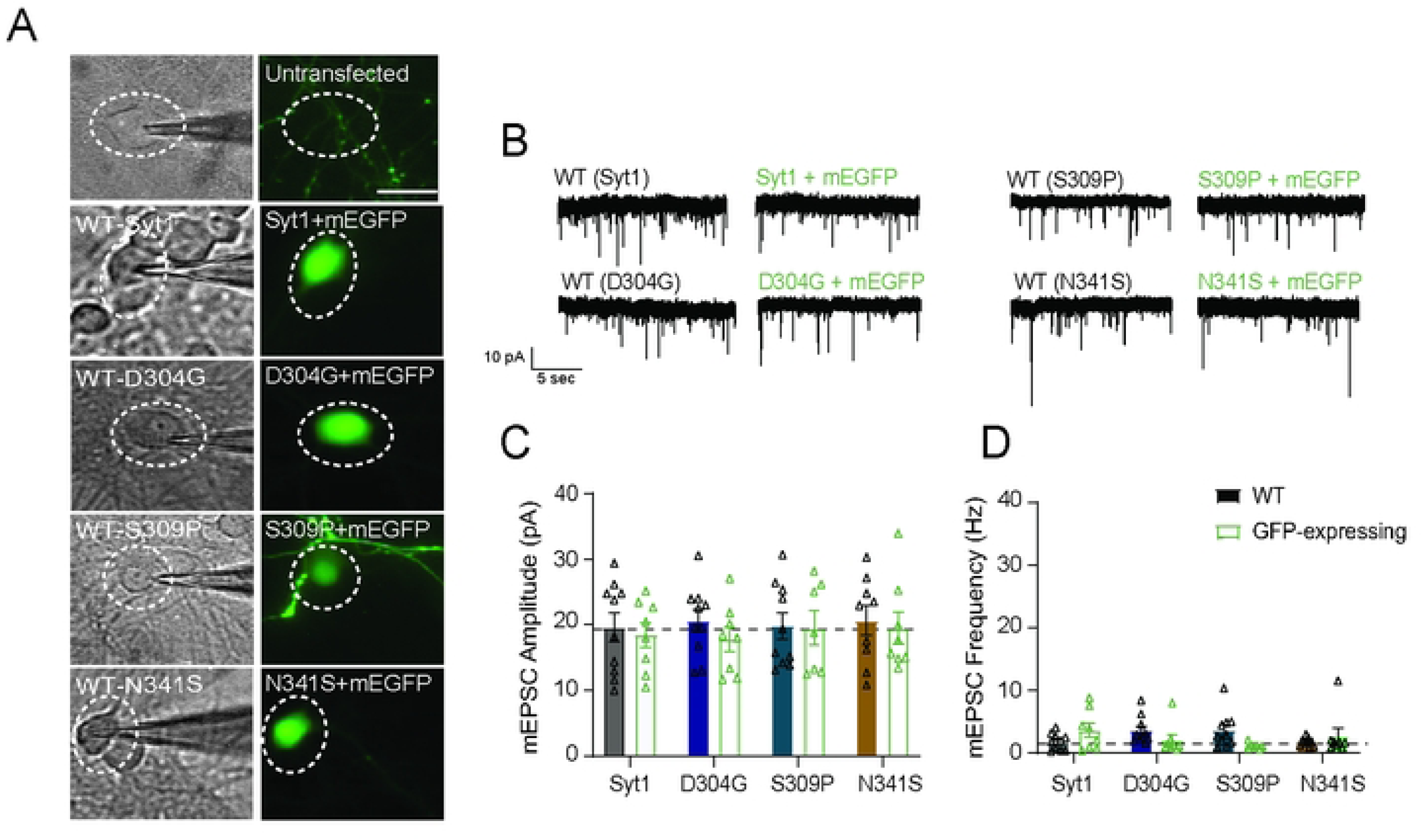
Cell autonomous expression of syt1 variants reveal presynaptic changes elicit chronic changes in spontaneous neurotransmission. (A) Representative image of networks expressing Syt1 variants and mEGFP in neurons. (B) Representative raw voltage clamp traces of WT un-transfected neurons and mEGFP-syt1 expressing neurons following sparse treatment. Scale bar, 50 µm. (C) Comparison of the mean mEPSC amplitude (pA) (Two-way ANOVA with Sidak’s multiple comparisons test: WT vs. Syt1+mEGFP p=0.991; WT vs. D304G+mEGFP p= 0.822; WT vs. S309P+mEGFP p>0.999; WT vs. N341S+mEGFP p=0.992; N=3, n=7-10 per group). (D) Comparison of the mean mEPSC frequency (Hz) (Two-way ANOVA with Sidak’s multiple comparisons test: WT vs. Syt1+mEGFP p=0.386; WT vs. D304G+mEGFP p= 0.542; WT vs. S309P+mEGFP p=0.141; WT vs. N341S+mEGFP p=0.884; N=3, n=7-10 per group).

### N341S-mutation occludes induction of homeostatic synaptic plasticity

Previous studies have elucidated how chronic signaling through spontaneous neurotransmission pathways elicits homeostatic synaptic plasticity, a well-studied compensatory mechanism in response to altered neuronal activity that allows neurons to fine-tune their signaling pathways, and ultimately, maintain stable circuitry and optimal network function^31–35^. Particularly, homeostatic plasticity is impaired in certain neurodevelopment disorders, making this process a compelling target for BAGOS pathophysiology^34^. Thus, we pharmacologically induced homeostatic plasticity pathways through both rapid and chronic signaling pathways. We acutely treated neuronal cultures expressing N341S or empty vector (CTRL) with dH_2_O as vehicle control, Tetrodotoxin (TTX; the voltage gated sodium channel blocker) alone, or TTX in combination with APV (N-methyl-D-Aspartate blocker) for 3 hours (Fig. 5A). In our CTRL neurons, we observed that 3-hour treatment of TTX+APV was sufficient to elicit rapid induction of homeostatic synaptic plasticity, illustrated by significantly increased mEPSC amplitudes with no change in frequency. Notably, our N341S expressing neuronal cultures failed to undergo rapid induction of homeostatic plasticity mechanisms, with no change in mEPSC amplitude or frequency (Fig.5 B, C).

**Figure 5:**
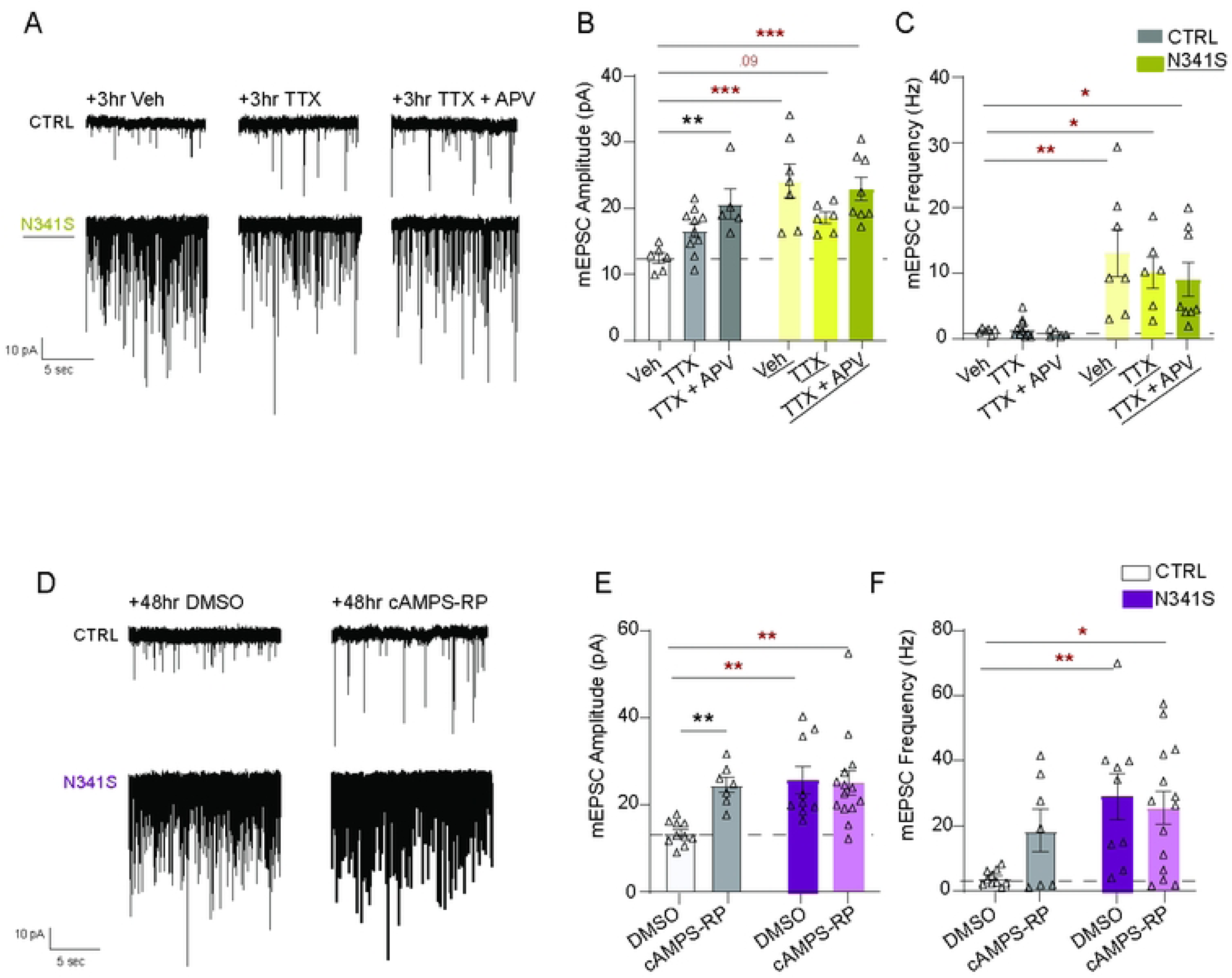
N341S-mutation occludes induction of homeostatic synaptic plasticity. (A) Representative raw traces of mEPSCs of Empty Vector (CTRL) and Syt1 variant (N341S) neurons following acute 1 hr treatment with vehicle (VEH), TTX, or TTX+AP5 treatment. (B) Comparison of mean baseline mEPSC amplitude (pA) for each group after incubations (Two-way ANOVA with Dunnett’s multiple comparisons test: VEH vs. TTX of CTRL p=0.120; VEH vs. TTX+APV of CTRL p=0.006; VEH vs. TTX of N341S p=0.050; VEH vs. TTX+APV of N341S p=0.796, N=2, n=6-7 per group). (C) Comparison of mean baseline mEPSC frequency (Hz) for each group after incubations (Two-way ANOVA with Dunnett’s multiple comparisons test: VEH vs. TTX of CTRL p=0.986; VEH vs. TTX+APV of CTRL p=0.991; VEH vs. TTX of N341S p=0.519; VEH vs. TTX+APV of N341S p=0.276, N=2, n=6-7 per group). (D) Representative raw traces of mEPSCs of empty vector (CTRL) and Syt1 variant (N341S) neurons after chronic 48 hr DMSO and cAMPS-RP treatment. (E) Comparison of mean baseline mEPSC amplitude (pA) for each group after incubations (Two-way ANOVA with Sidak’s multiple comparisons test: DMSO vs. CAMPS-RP of CTRL p=0.007; DMSO vs. cAMPS-RP of N341S p=0.865; N=2-3, n=7-14 per group). (F) Comparison of mean baseline mEPSC frequency (Hz) for each group after incubations (Two-way ANOVA with Sidak’s multiple comparisons test: DMSO vs. CAMPS-RP of CTRL p=0.170; DMSO vs. cAMPS-RP of N341S p=0.856; N=2-3, n=7-14 per group).

Previous work from our lab has demonstrated an activity-independent approach to induce homeostatic synaptic plasticity using chronic blockade of cAMP signaling^36^. Here, we utilized this approach and chronically treated neuronal cultures for 48 hr with either DMSO vehicle control, or cAMPS-RP (cAMP inhibitor) (Fig. 5D). CTRL neurons showed robust homeostatic synaptic upscaling; however, the N341S expressing cultures fail to augment mEPSC amplitudes after treatment (Fig. 5E, F). These data demonstrate that the N341S-disease mutation alone augmented mEPSC amplitudes 1-2 fold at baseline, but occluded any subsequent action of homeostatic upscaling mechanisms on post-synaptic signaling pathways.

### Substituting N341 residue with neutral amino acids blocks spontaneous release dysregulation seen with the serine mutation

Predictive modeling of this particular N341S syt1 mutation was suggested it to be “possibly damaging”; however, no functional evidence has been provided to demonstrate this premise^24^. We sought to determine whether this serine mutation alone is sufficient to affect syt1 function, or whether other mutations at the same residue replicate the severe deficits in spontaneous release we observed. We created artificial mutants by substituting isoleucine or alanine to test whether introducing nonpolar residues recapitulates the effects of mutating this site, and more broadly, whether this residue is sensitive to any mutation. Because serine can be phosphorylated, we also generated a negatively charged phosphomimetic mutant by substituting aspartic acid for the serine (Fig. 6A). Unlike the N341S, when we expressed 3 of the artificial mutants (N341I, N341A, or N341D), they demonstrated similar levels of spontaneous neurotransmission to that of CTRL (Fig. 6B, C). We do not observe any changes with N341D mutation, suggesting that substantial augmentation of spontaneous neurotransmission is specific to serine at this position.

**Figure 6:**
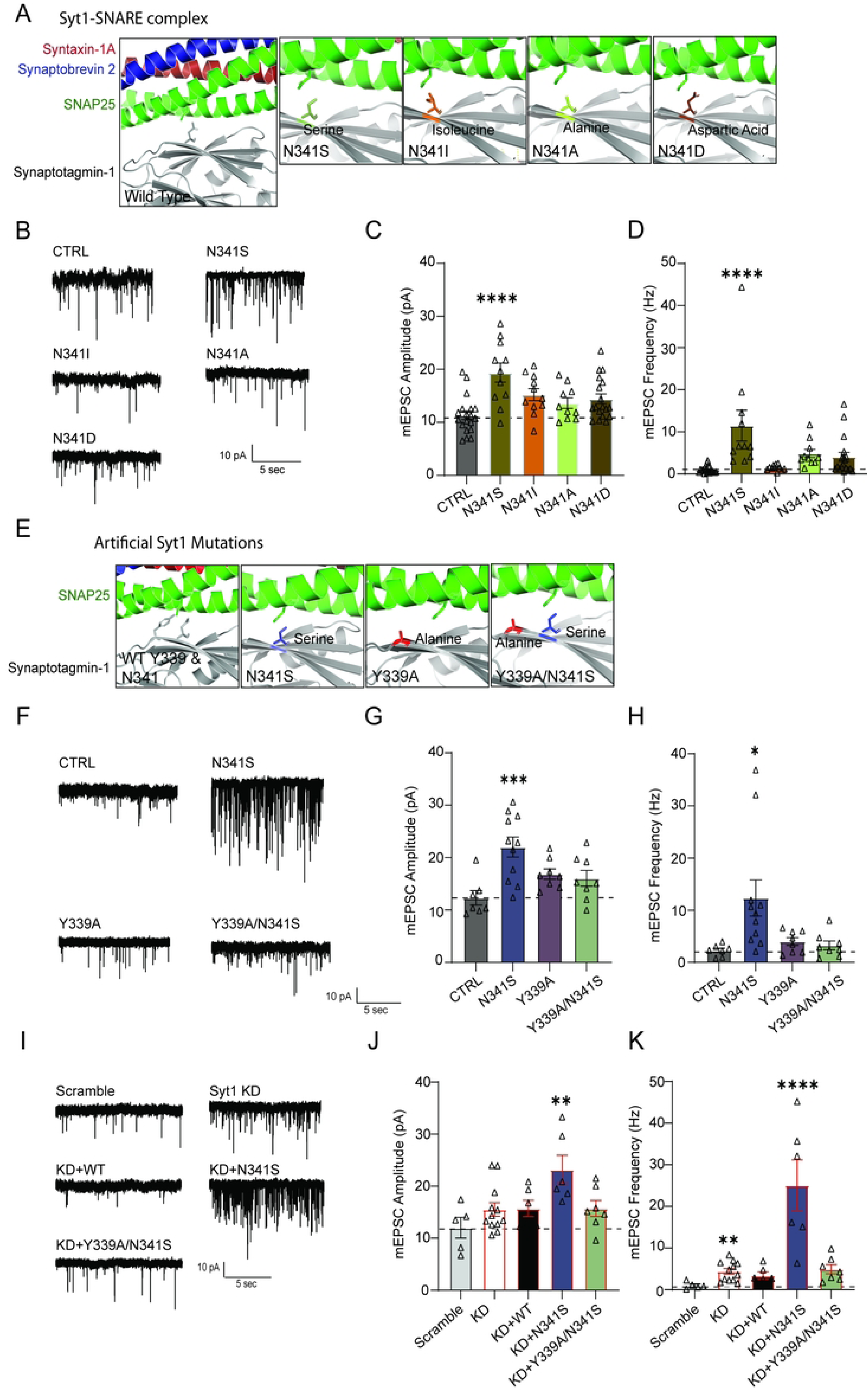
Substituting N341 residue with neutral amino acids blocks spontaneous release dysregulation seen with the serine mutation. (A) Zoomed in rendering of Syt1-SNARE complex structure, with panels illustrating artificial point mutations at N341 residue in syt1 protein. Crystal structure rendering modified using PYMOL (PDB: 5CCG). (B) Representative raw traces of baseline mEPSCs on empty vector (CTRL), disease variant (N341S), and other N341 mutations (N341I, N341A, N341D). (C) Comparison of baseline mEPSC amplitudes (pA) (One-way ANOVA with Dunnett’s multiple comparisons test: CTRL vs. N341S p<0.0001; CTRL vs. N341I p=0.050; CTRL vs. N341A p=0.435; CTRL vs. N341D p=0.083; N=2-6, n=10-19 per group). (D) Comparison of mEPSC frequencies (Hz) (One-way ANOVA with Dunnett’s multiple comparisons test: CTRL vs. N341S p<0.0001; CTRL vs. N341I p=0.998; CTRL vs. N341A p=0.241; CTRL vs. N341D p=0.299; N=2-6, n=10-19 per group). (E) Zoomed in rendering of syt1 N341 residue with Y339 and N341 residues highlighted in gray in same panel. Additional panels showing artificial point mutation Y339A, patient N341S mutation pictured in another panel, and other mutations together in same panel. (F) Representative raw traces of baseline mEPSCs on CTRL, N341S, Y339A, and double mutant Y339A/N341S. (G) Comparison of baseline mEPSC amplitudes (pA) for each group (One-way ANOVA with Dunnett’s multiple comparisons test: CTRL vs. N341S p=0.0005; CTRL vs. Y339A p=0.163; CTRL vs. Y339A/N341S p=0.291; N=3-5, n=7-11 per group). (H) Comparison of mEPSC frequencies (Hz) for each group (One-way ANOVA with Dunnett’s multiple comparisons test CTRL vs. N341S p=0.012; CTRL vs. Y339A p=0.925; CTRL vs. Y339A/N341S p=0.983; N=3-5, n=7-11 per group). (I) Representative raw traces of baseline mEPSCs on scramble (SCR), Syt1 knockdown (KD), knockdown with wildtype rescue (KD+WT), knockdown with N341S rescue (KD+N341S), and knockdown with rescue of double mutant (KD+Y339A/N341S). (J) Comparison of baseline mEPSC amplitudes (pA) for each group (One-way ANOVA with Dunnett’s multiple comparisons test: SCR vs. KD p=0.491; SCR vs. KD+WT p=0.560; SCR vs. KD+N341S p=0.002; SCR vs. KD+Y339A/N341S p=0.532, N=2-4, n=5-12 per group). (K) Comparison of mEPSC frequencies (Hz) for each group (One-way ANOVA with Dunn’s multiple comparisons test: SCR vs. KD p=0.009; SCR vs. KD+WT p=0.606; SCR vs. KD+N341S p<0.0001; SCR vs. KD+Y339A/N341S p=0.131; N=2-4, n=5-12 per group).

### Simultaneously mutating 339 and 341 residues reverses upscaling seen with ser341 mutation

Previous studies with the SNARE complex have demonstrated that syt1 and SNAP-25 interact at primary, secondary, and tertiary interfaces, cooperating in a manner that promotes vesicle fusion^37^. The largest region that forms the primary interface occurs between the C2B domain of syt1 and SNAP25, and harbors critical residues for functional syt1/SNARE complex binding^38^. Knowing this patient-relevant serine mutation lies nearby a critical residue in the primary interface (Y339), we wanted to test whether mutating this amino acid mitigates, or possibly replicates, what we observe with N341S (Fig. 6D). We tested the following three mutations: an artificial syt1 mutation Y339A, a Y339A/N341S double mutation, and the N341S patient-relevant mutation. Strikingly, the single Y339A mutation alone did not augment spontaneous neurotransmission. Uniquely, we did find that the Y339A/N341S double mutation rescued aberrant spontaneous neurotransmission back to CTRL levels (Fig. 6E,F).

To further assess whether this double mutation can maintain spontaneous neurotransmission like wild type syt1, we recreated the syt LOF background using rat syt1 shRNA and performed rescue experiments using these mutations. The double mutation Y339A/N341S mirrored the same phenotype, with spontaneous neurotransmission rescued in the scramble (negative control) condition (Fig. 6G, I). These results demonstrate that this serine mutation at residue 341 is sufficient to disrupt intermolecular interactions of syt1/SNAP-25 at the primary interface, and chronically augment spontaneous neurotransmission across both frequency and amplitude.

### Recombinant GST-HA-Syt1C2A/B-N341S protein is phosphorylated

To determine whether the serine mutation disrupts spontaneous release by introducing a novel phosphorylation site, we examined recombinant GST-HA-syt1 protein phosphorylation in brain homogenate using a phospho-serine antibody and GST-pull-down assays. We conducted GST-pulldowns from brain lysate to assess whether WT syt1 and the patient variant N341S-syt1 is phosphorylated (Fig. 7A). We further tested whether exposure to different buffer conditions alters the phosphorylation readout; notably, N341S-syt1 exhibited a significantly higher phospho-serine signal than WT-syt1 across all buffer conditions (Fig. 7B). A limitation of this approach is that many proteins have phospho-serine residues and the affinity of isolated phospho-serine residue to the antibody is not strong enough to generate clean signals, which likely accounts for the low-level of background signal observed in the WT-syt1 condition. We also found that recombinant syt1-N341S protein generated in *E. coli* shows a significantly higher phospho-serine signal than WT, with at least a threefold increase across all protein loading amounts (Fig. 7C,D) consistent with the premise that this mutation generates a phosphorylation site.

**Figure 7:**
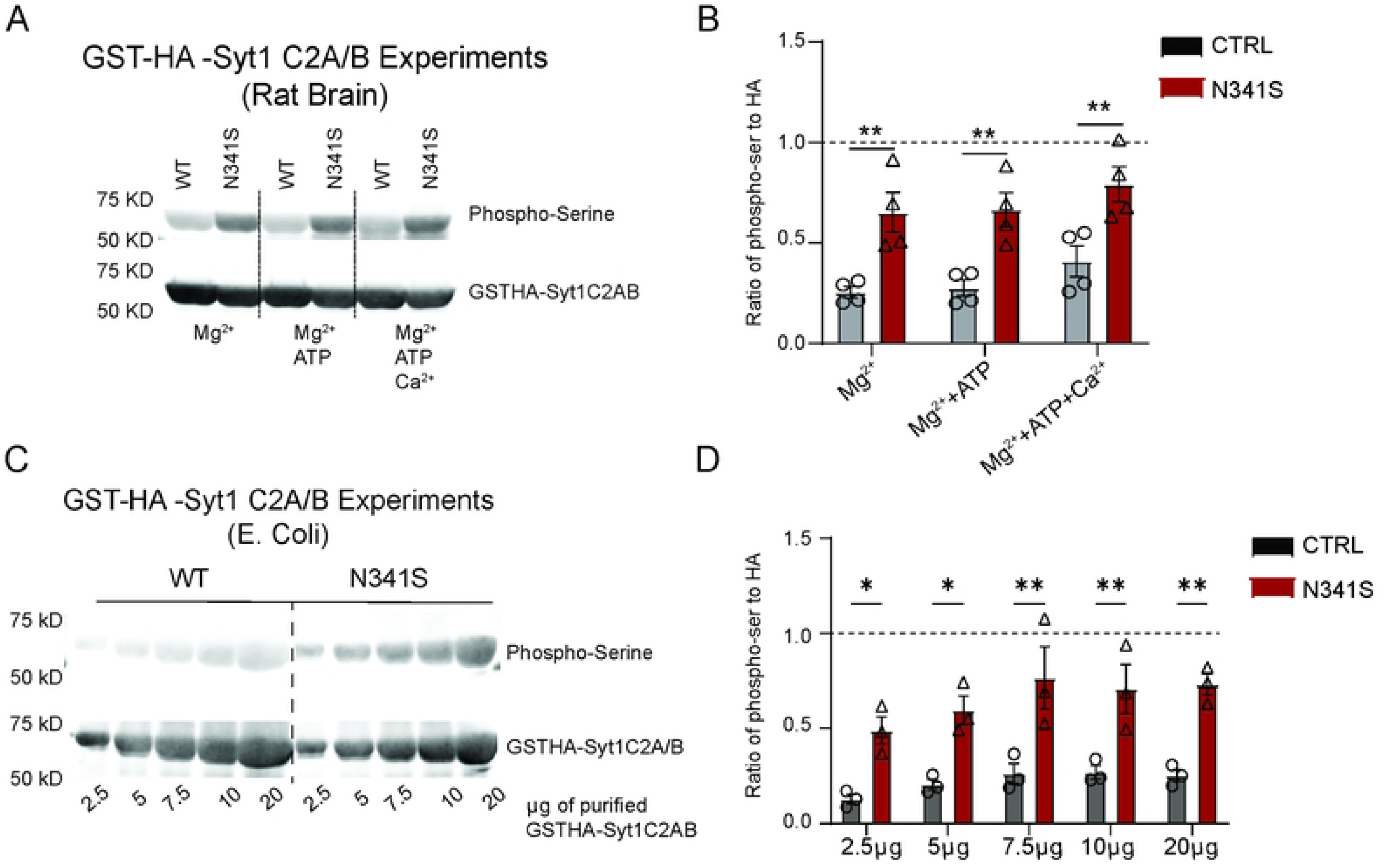
N341S protein is phosphorylated following GST-pulldowns. (A) Immunoblot image of pGEX-HA-Syt1C2A/B or pGEX-HA-Syt1C2A/B-N341S. Recombinant pGEX-HA-Syt1 (WT or patient variant N341S) protein was added to rat brain homogenate and pulled down using GST beads. Proteins were quantified and analyzed by SDS-PAGE and immunoblotting to detect anti-phospho-serine and HA-tag. (B) Comparison of phospho-serine protein normalized to HA-tag signal for each group (Two-way ANOVA with Sidak’s multiple comparisons test: [for Mg^2+^] WT vs. N341S p=0.003; [for Mg^2+^+ATP] WT vs. N341S p=0.004; [for Mg^2+^+ATP+Ca^2+^] WT vs. N341S p=0.005; n=3). (C) Immunoblot image of GST-HA-Syt1 generated in *E. coli.* Recombinant GST-HA-Syt1 protein was generated in E. coli and purified proteins were quantified and analyzed by SDS-PAGE and immunoblotting. Immunoblot protocol is the same (as detailed in Methods). (D) Proteins were normalized to HA- tag signal for each group (Two-way ANOVA with Sidak’s multiple comparisons test: [for 2.5µg] WT vs. N341S p=0.021; [for 5µg] WT vs. N341S p=0.011; [for 7µg] WT vs. N341S p=0.001; [for 10µg] WT vs. N341S p=0.004; [for 20µg] WT vs. N341S p=0.001; n=3).

### Inhibiting kinase activity rescues functional spontaneous neurotransmission in ser341 mutation

So far, we found that the ser341 mutation lies adjacent to the primary interface, and that introducing an alanine near this serine restores the augmented spontaneous release phenotype down to wild type levels. Our next experiment aimed to address whether introducing a serine at residue 341 next to the endogenous tyrosine at residue 339 creates a phosphorylation site. Here, we built on previous findings from our group on a patient-relevant SNAP-25 mutation in the syt1/SNAP-25 primary interface (SNAP-25 D166Y), which produced a similar phenotype of increased mEPSC amplitude and frequency compared to WT^39^. The SNAP25 mutation introduces a tyrosine at residue 166, creating a new potential phosphorylation site that may contribute to the disease phenotype. To test whether phosphorylation occurs as a consequence of these mutations, we acutely treated CTRL, syt1-N341S, or SNAP-25-D166Y variant expressing neurons with staurosporine (a wide spectrum serine/threonine kinase inhibitor) for 1 hr (Fig. 8A). We observed that acute 1hr treatment with this broad kinase inhibitor significantly decreased mEPSC frequency in both the syt1-N341S and the SNAP-25-D166Y variant, but did not affect mEPSC amplitudes (Fig. 8B, C). Acute treatment had no effect on mEPSC amplitude or frequency in the CTRL as well, but kinase inhibition strikingly restored mEPSC frequency closer to CTRL levels despite recordings being performed in inhibitor-free conditions. We also tested chronic treatment of syt1-N341S expressing neurons with selective kinase inhibitors, such as U0126 and Trametinib (MEKI/II kinase inhibitors), and did not observe any effect (See Supp. 4.)

**Figure 8:**
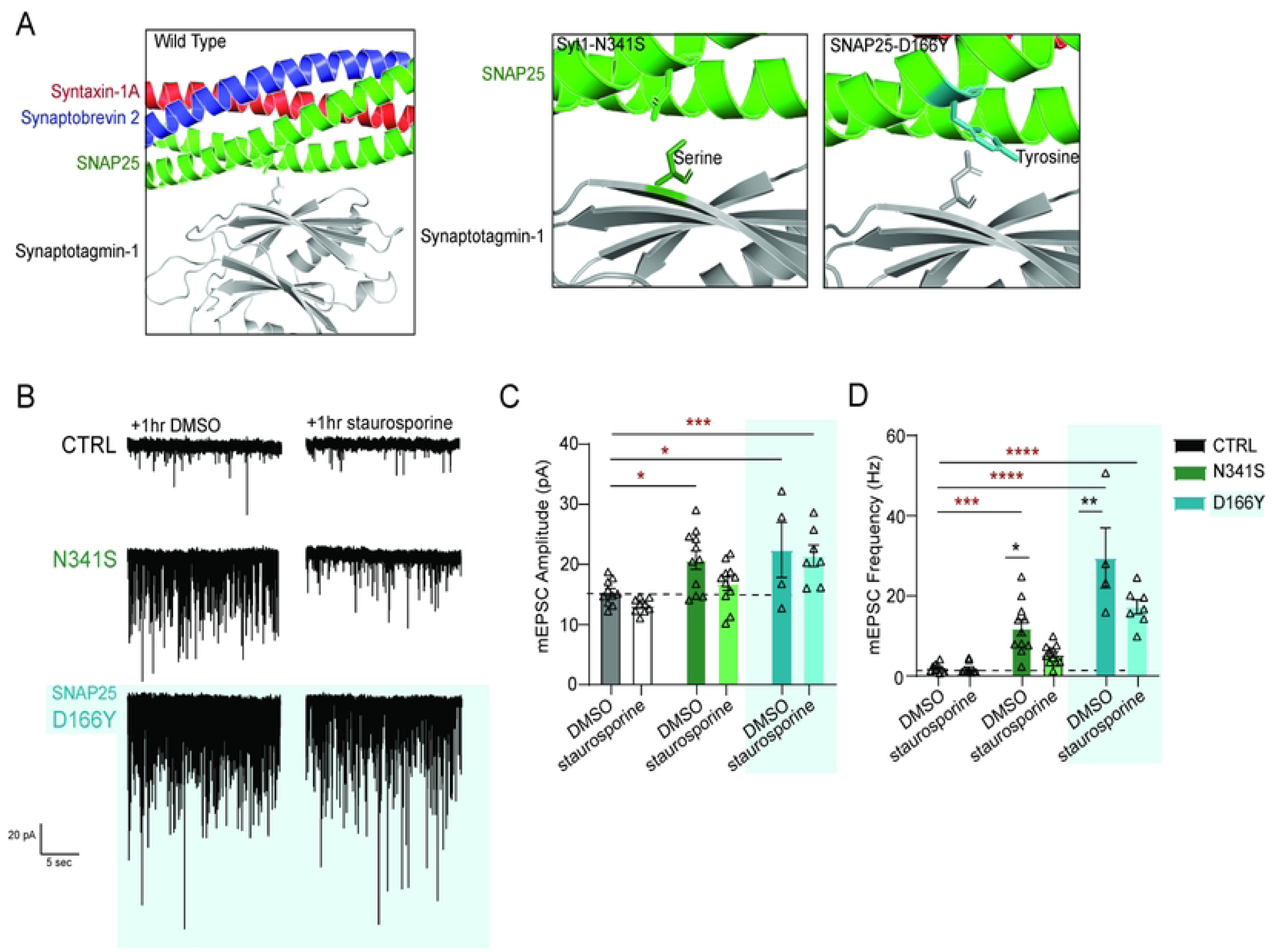
Inhibiting kinase activity rescues functional spontaneous neurotransmission back to wild-type phenotype. (A) Zoomed in rendering of Syt1-SNARE complex primary interface structure, with panels illustrating patient mutations in syt1 and SNAP25 that sit in interface. Crystal structure rendering modified using PYMOL (PDB: 5CCG). (B) Representative raw traces of mEPSCs of empty vector (CTRL), syt1-mutated disease variant (N341S), and SNAP25-mutated disease variant (D166Y) neurons at baseline and after acute 1 hr treatment with 1 µM staurosporine. (C) Comparison of mean baseline mEPSC amplitude (pA) for each group (Two-way ANOVA with Sidak’s multiple comparisons test: CTRL vs. N341S [DMSO] p=0.010; CTRL vs. D166Y [DMSO] p=0.013; CTRL vs. N341S [staurosporine] p=0.121; CTRL vs. D166Y [staurosporine] p=0.0006, N=1-2, n=10-11 per group.) (Two-way ANOVA with Sidak’s multiple comparisons test [before versus after staurosporine]: [CTRL] DMSO vs. staurosporine p=0.615; [N341S] DMSO vs. staurosporine neurons p=0.098; [D166Y] DMSO vs. staurosporine p=0.976, N=1-2, n=10-11 per group). (D) Comparison of mean baseline mEPSC frequency (Hz) for each group (Two-way ANOVA with Sidak’s multiple comparisons test: CTRL vs. N341S [DMSO] p=0.0002; CTRL vs. D166Y [DMSO] p<0.0001; CTRL vs. N341S [staurosporine] p=0.283; CTRL vs. D166Y [staurosporine] p<0.0001, N=1-2, n=10-11 per group). (Two-way ANOVA with Sidak’s multiple comparisons test [before versus after staurosporine]: [CTRL] DMSO vs. staurosporine p=0.999; [N341S] DMSO vs. staurosporine p=0.016; [D166Y] DMSO vs. staurosporine p=0.002, N=1-2, n=10-11 per group)

## DISCUSSION

### Summary of Results

Although BAGOS is an extremely rare disease, studying pathogenic mutations in conserved machinery provides a framework for understanding how a single protein mutation can affect both neurotransmission and synaptic plasticity. Poor gene tolerance, which indicates high vulnerability to mutations, is observed across canonical SNARE proteins and their regulatory proteins, such as syt1^40^. Because the structure of the SNARE complex and its associated proteins is highly conserved across eukaryotes, even subtle shifts in assembly can precipitate structural and functional changes, driving disease.

Currently, there are 30 identified pathogenic variants in synaptotagmin-1 protein, with patients presenting a spectrum of phenotypes of BAGOS Disorder (i.e. Syt1-associated neurodevelopmental disorder)^6,24,25,41^. *In vitro* analyses have revealed disrupted evoked release, most notably a loss of synchronous release accompanied by elevated spontaneous neurotransmission^6^. Functional characterization has been performed on 5 of the syt1 *de novo* variants, expressed in cortical *SYT1* KO neurons, which demonstrated a range of phenotypes and graded deficits in evoked release^56^. Several variants also displayed impaired calcium-binding, consistent with syt1’s role as a Ca^2+^-sensor in vesicle fusion^17,42–44^. Although many variants remain uncharacterized at the molecular level, recent computational modeling has provided valuable predictions regarding their potential effects^6^, which include perturbations in cation binding, altered protein-protein interactions, and changes to surface charge.

Our study explores how specific syt1 variants contribute to the underlying disease mechanisms and to the variability in clinical presentation observed among patients. Our results show that syt1 plays a key role in Ca^2+^-dependent vesicle exocytosis at both excitatory and inhibitory synapses, and that patient-relevant syt1-variants can alter this process. We examined 11 missense mutations in syt1 and found that over-expressing three C2B variants—D304G, S309P, and N341S—significantly impacted spontaneous excitatory neurotransmission, with N341S also augmenting spontaneous inhibitory neurotransmission (Fig. 1). When expressed on a syt1-KD background, these variants produced even greater effects, with N341S driving spontaneous release beyond the increase seen by syt1-KD alone (Fig. 3).

In experiments where we probed evoked neurotransmission with variants expressed on wild-type networks, we did not observe any significant disruption across all C2A and C2B variants (Fig. 2). Surprisingly, we did not detect any significant difference in evoked release across D304G, S309P and N341S variant networks despite seeing significant changes to both excitatory and inhibitory spontaneous neurotransmission. Conversely, previous work has shown that rescuing SYT1 KO synapses with D304G mutation reduces evoked neurotransmission ^29,49^. This discrepancy can perhaps be attributed to the nature of lentiviral expression compared to rescue of SYT1 KO networks consistent with dose-dependent, graded potency of the syt1-patient variants^5^. Differences in our findings may also be attributed to a three-fold difference in baseline evoked responses observed in wild-type conditions between autaptic and high-density culture systems^49^.

We found that the enhanced spontaneous release phenotypes originate from presynaptic changes, as sparse networks containing both syt1-variant and WT neurons displayed normal activity, suggesting that syt1 variants disrupt wild-type function when globally expressed rather than through single- synapse alterations (Fig. 3). We showed that the N341S variant failed to induce homeostatic synaptic plasticity under both rapid and chronic induction conditions, suggesting that this mutation disrupts compensatory pathways required to maintain stable network activity (Fig. 4). However, we cannot exclude the possibility that these networks have already engaged homeostatic mechanisms. Consequently, our induction paradigm may be limited in its ability to determine whether a homeostatic threshold has already been reached, or whether these mechanisms are themselves insufficient to reveal which regulatory pathways remain intact. Despite dysregulated spontaneous neurotransmission in N341S networks, evoked transmission remained largely intact. Nevertheless, because homeostatic plasticity depends on spontaneous neurotransmission, these findings suggest impaired coordination of spontaneous signaling necessary for effective homeostatic regulation.

Given these disruptions in distinct modes of neurotransmission and plasticity, we asked whether the serine substitution at residue 341 directly drives these downstream effects, or whether nearby residues also have the capacity to modulate this outcome. Mutating residue 341 to neutral amino acids (N341A and N341I) reversed the N341S loss-of-function phenotype, highlighting that a change from an asparagine to a neutral amino acid is insufficient for eliciting changes in spontaneous neurotransmission.

Is the asparagine to serine introducing a novel phosphorylation site? To test this region’s sensitivity further, we introduced a second mutation at a tyrosine residue two amino acids upstream of the serine (Y339A/N341S), which restored spontaneous release to wild-type levels. We observed that mutating the tyrosine alongside the intact serine was sufficient to functionally rescue neurotransmission back to WT levels. Taken together, our results highlight how this region of the C2B domain is highly sensitive to specific permutations at the primary interface, ultimately highlighting the pathogenic effects triggered by introducing a serine at residue 341 near the primary syt1/SNAP-25 interface.

Finally, treatment with a broad serine/threonine kinase inhibitor for 1hr rescued the elevated spontaneous release observed in both syt1-N341S and SNAP-25-D166Y mutants, implicating phosphorylation at this interface as a potential mechanism driving hyperactive spontaneous neurotransmission.

### General Implications on Therapeutic Findings

Given the efficacy of a broad kinase inhibitor, we next asked whether phosphorylation arises specifically from the serine substitution at position N341. To test this, we generated recombinant syt-N341S protein to assess whether the mutation introduces a novel phosphorylation site. Using a phospho-serine antibody, we found that N341S exhibited significantly elevated serine phosphorylation compared to WT, regardless of whether the protein was incubated with brain homogenate or expressed in *E. coli* (Fig. 7). Together these findings suggest that phosphorylation—or other post-translational modifications—at the syt1-SNARE interface may disrupt complex interactions and consequently enhance spontaneous neurotransmission.

While the syt1–SNARE interface is vital for proper complex folding and interaction, the impact of disease-associated mutations remains under-studied. Here, we demonstrate that two distinct mutations at this interface produce similar phenotypes rescued by kinase inhibition, suggesting that phosphorylation-dependent disruption of the SNARE machinery is a common pathophysiological trigger. Syt1-associated disorders provide a framework for understanding how the selective dysregulation of specific release modes can alter signaling without impairing all forms of neurotransmission. This molecular convergence highlights the potential for shared therapeutic targets across clinically heterogeneous SNAREopathies^5,47^.

Beyond neurotransmission, future research must address how mutations like N341S affect vesicle priming and SNARE disassembly. Identifying these specific cellular markers will improve our ability to predict clinical severity based on variant location. Ultimately, parsing the molecular nuances of these overlapping disorders will guide the development of targeted therapeutics to restore functional neurotransmission and plasticity.

## Author Contributions

E.D.B. conducted electrophysiology, immunoblotting analysis, and fluorescence microscopy experiments. O.S. designed lentiviral constructs and performed GST-pulldown assays and immunoblotting. Q.Z. designed lentiviral constructs. E.D.B. and R.M.A. conducted formal analysis. E.D.B. and E.T.K. contributed to the initial conceptualization of the study. E.T.K. supervised the project and contributed to its overall design and direction. E.D.B. and E.T.K. wrote the manuscript.

## MATERIALS AND METHODS

### Animals

Pregnant Sprague-Dawley rats (Charles River) were housed individually until they gave birth to a litter. Sprague-Dawley rat pups of either sex were used for the rat hippocampal cultures. All animal procedures were performed in accordance with the guide for the care and use of laboratory animals and were approved by the Institutional Animal Care and Use Committee at Vanderbilt University. Health status of the live animals was periodically checked and confirmed by the veterinary staff of animal facilities of Vanderbilt University.

### Primary hippocampal culture preparation

Hippocampal rat cultures were generated from postnatal (P) day 1/2 wildtype pups from either sex per standard protocols^11^. Pups were rapidly decapitated and hippocampi (WT P1/2 pups) were isolated from each hemisphere. Hippocampi were dissected in 20% FBS (fetal bovine serum) containing Hank’s balanced salt solution, and then washed and treated with 10 mg/mL trypsin and 0.5 mg/mL DNAse at 37°C for 10 min. Tissue was washed again, dissociated with a P1000 tip, and centrifuged at 1200 rpm for 10 min at 4°C. Matrigel coated coverslips were prepared as previously described^11^. Cells were then resuspended and plated on Matrigel-coated glass coverslips in 24-well plates at a density of six coverslips per hippocampus. Cultures were kept in humidified incubators at 37°C for and gassed with 95% air and 5% CO_2_. Plating media contained 10% FBS, and 20 mg/L insulin, 2 mM L-glutamine, 0.1 g/L transferrin, 5 g/L D-glucose, 0.2 g/L NaHC03 in minimal essential medium (MEM). After 24 hours (hr), plating media was exchanged for growth media containing 4 μM cytosine arabinoside (as well as 5% FBS, 0.5 mM L-glutamine, and B27) to inhibit glial proliferation. On day *in vitro* (DIV) 4, growth media was exchanged to a final concentration of 2 μM cytosine arabinoside. For lentiviral gene expression, neurons were infected with the lentivirus containing our plasmid of interest at DIV 4. For neurons that underwent sparse transfection experiments, transfection was performed on DIV 7 (see below).

### Cell lines

Human embryonic kidney-293 (HEK293-T) cells were used to generate lentiviruses for neuronal infection. HEK293-T cells were maintained in Dulbecco’s Modified Eagle Medium supplemented with 10% FBS, penicillin, and streptavidin in incubators held at 5% CO_2_ at 37°C. Cells were passaged and split at ∼75% confluency.

### Sparse neuron transfection

Neuronal sparse transfections were performed on DIV 7 using a Ca^2+^ phosphate kit (ProFection Mammalian Transfection System, Cat #E1200, Promega), as previously described^12^. A DNA/calcium phosphate mixture was prepared as follows (per well/24-well plate): 1 µg of plasmid DNA, 2 µL of 2 M CaCl_2_, and 15 µL dH_2_O. This mixture was then added dropwise to an equal volume of 23 N-2-Hydroxyethylpiperazine-N0 -2-Ethanesulfonic Acid (HEPES), while constantly low-vortexing between drop additions. The combined solution was allowed to form for 15 minutes at room temperature. Neuron conditioned media (1 mL/well) was saved and replaced with warmed MEM, and 30 µL of plasmid mixture was added dropwise to each well. Plates were returned to 5% CO_2_ incubator at 37°C for 30 minutes, followed by washing the cells twice with 1 mL MEM, after which previously saved conditioned media was added back to each well. Neurons were used for experiments at DIV 15–18. Transfected neurons could be visualized by their GFP expression from the transfected construct starting 3 days after transfection.

### Lentiviral preparation

Lentiviruses were prepared by transfection of HEK293-T cells using Fugene 6 (Roche) with the plasmid of interest at 1.0 µg, together with three packaging plasmids at 0.5 µg each (pRSV-RCTRL, pCMV-VSV-G, and pMDLg/pRRE) (Addgene). Culture medium on the cells was exchanged for neuronal growth media 24 hr after transfection. The supernatant of neuronal growth media containing viruses were harvested 3 days after transfection and applied to the primary hippocampal cultures. Using this approach, we consistently obtained infection frequencies approaching ∼100%.

### Drug treatment

For rapid homeostatic plasticity experiments, neurons were incubated for 3 hours with either 1 µM Tetrodotoxin (TTX) alone (TOCRIS), 50 µM AP5 alone (TOCRIS), or both combined. To model homeostatic plasticity using chronic pharmacological effect, neurons were incubated for 48 hours with cAMPS-RP (MedChemExpress). DMSO (Sigma-Aldrich) and H_2_O were used as vehicle controls. To inhibit serine/threonine kinase function, neurons were acutely treated for 1 hour with 1 µM staurosporine (Sigma-Aldrich). Supplement experiments treated neurons for 1 hour with either 1 µM Trametinib (MedChemExpress) or 10 µM U0126 (Selleck Chem) as selective MEK I/II inhibitors.

### Electrophysiology

Whole-cell patch clamp recordings were performed on pyramidal neurons at DIV14-21 at a clamped voltage of -70 mV. Recordings were 3-5 minutes in length. All recordings were performed using a Burleigh PCS-5000 headstage, Axopatch 200B amplifier, Digidata 1550B digitizer, and Clampex 11.1 software (Molecular Devices). Internal pipette solution contained 115 mM CsMeSO3, 10 mM CsCl, 5 mM NaCl, 10 mM HEPES, 0.6 mM EGTA, 20 mM tetraethylammonium chloride, 4 mM Mg-ATP, 0.3 mM Na_3_GTP, pH 7.3-7.3, and10 mM QX-314 [N-(2,6-dimethylphenyl-carbamoylmethyl)-triethylammonium bromide]. The extracellular Tyrode’s solution contained 150 mM NaCl, 4 mM KCl, 2 mM CaCl_2_, 10 mM glucose, and 10 mM HEPES (pH7.4), 308 mOsm. To isolate AMPA-mediated mEPSCs, 1 µM TTX, 50 µM AP-5 and 50 µM PTX (or 50 µM bicuculline) were included in the bath. To isolate excitatory eIPSCs, APV (50μM) and CNQX (10 μM) were added to the bath. A parallel bipolar electrode provided field stimulation at 35 mA for evoked recordings.

### Determination of Syt1-N341S phosphorylation using rat brain homogenate

Recombinant GST-HA-Syt1C2A/B proteins were generated in BL21-CodonPlus (DE3)-RIPL chemical competent E. coli (Cat# 230280; Agilent) after transformation with either pGEX-HA-Syt1C2A/B or pGEX-HA-Syt1C2A/B-N341S. Purified GST-HA-Syt1C2A/B protein was immobilized to Glutathione Sepharose beads (Cytiva) by 1 hr rocking at 4°C. Beads were washed (3x) using a Tris buffer (20mM Tris pH 7.5, 100mM NaCl, 1mM EDTA, and 1mM EGTA). Washed beads were mixed with rat brain homogenate in either 1 mL of Mg^2+^ buffer, Mg^2+^ + ATP buffer, or Mg^2+^ +ATP+ Ca^2+^ buffer. Sample was incubated with inverted mixing for 1hr at 30°C. Beads were washed (5x) with corresponding buffers (Mg^2+^ buffer, Mg^2+^ + ATP buffer, or Mg^2+^ +ATP+ Ca^2+^ buffer) without brain homogenate. Proteins were quantified with SDS-PAGE and immunoblotting (as detailed above). Proteins were normalized to HA-tagging signal. Antibodies used were Anti-Phosphoserine (p5) (Rabbit) antibody (Rockland) and HA-tag Monoclonal antibody (Invitrogen).

### Production of recombinant GST-HA-Syt1C2A/B proteins in E. coli

For the generation of recombinant protein using E. coli, 20 µL of BL21-CodonPlus (DE3)-RIPL chemical competent E coli (from Agilent Cat# 230280) is transformed by adding 1 µL of ∼300 ng/ µL pGEX-HA-Syt1C2A/B or pGEX-HA-Syt1C2A/B-N341S constructs by a chemical method. Briefly, mixed plasmid and E. coli, incubated on ice for 30 min, heat shocked 42°C for 30 sec, placed on ice for 2 min, added 1 ml LB medium, incubated 37°C for 1 hr, spun down at 9000 rpm for 5 min using a microcentrifuge, discarded ∼900 µL supernatant, resuspended E. coli using ∼100 µL leftover supernatant, and spread E. coli on a LB agar plate (10g/L Tryptone, 5g/L Yeast extract, 5g/L NaCl, 2% agar) containing 50 µg ampicillin/ml LB). Proteins were quantified and analyzed with SDS-PAGE and immunoblotting (as detailed above). Proteins were normalized to HA-tagging signal. Antibodies used were Anti-Phosphoserine and HA-tag Monoclonal antibody (same as detailed above).

### SDS-PAGE

Samples were loaded into a 10% sodium dodecyl sulfate-polyacrylamide gel electrophoresis (SDS-PAGE) gel. Protein was transferred with a Trans-Blot Turbo Transfer System to a PVDF membrane. Membranes were then blocked in 5% bovine serum albumin (A1470-25G, Sigma) for 45 minutes, and then incubated overnight with primary antibodies in 5% milk at 4°C at the following dilutions: mouse anti-Phosphoserine (Invitrogen), rabbit anti HA-tag (Rockland). After incubation, membranes were washed 3 times in TBS for 5 minutes. Membranes were then incubated for 1 hr at with secondary antibodies at the following dilutions (1:10,000): goat anti-rabbit IgG secondary antibody (IRDye® 800CW, Li-COR), donkey anti-rabbit secondary antibody (IRDye® 680LT,Li-COR). Membranes were imaged with Odyssey Clix imaging machine (Li-COR). Phospho-serine band intensities were calculated using Fiji ImageJ and normalized to their respective HA-tag.

### Quantification and Statistical Analysis

Mini Analysis software (Synaptosoft) was used to calculate mPSC amplitudes and frequencies. Clampfit 11.1 (Molecular Devices) was used to analyze ePSCs. Statistical analyses were performed using Prism 10.6.1 (GraphPad Software). All data are reported as mean ± SEM. An unpaired two-tailed t test was used for comparison when comparing two groups. For parametric analysis of multiple comparisons, two-way analysis of variance (one-way ANOVA and two-way ANOVA) with Tukey post hoc or Dunnett’s test were used. For other data, a Kruskal-Wallis test followed by Tukey’s post hoc correction was used for multiple comparisons. Outliers were identified with Robust regression and Outlier removal (ROUT) method. Differences among experimental groups were considered statistically significant when a p value < 0.5 was reached. Significant symbols used in figures are defined as such: *p<0.05; **p<0.01; ***p<0.001;****p<0.0001; n.s., not significant. The number of neurons for each experiment are given in the figure legends (N=biological replicates, n=technical replicates), and individual data points represent neurons, unless indicated otherwise.

## Acknowledgements

We would like to thank Clara McCarthy, Abigael Weit, Alp Uzay and Christine Saunders for their insightful comments on the manuscript. This work was supported by the National Institute of Neurological Disorders and Stroke (grant R01NS134128-23.) Figures 1A,1B, 2A, and 3A were created with BioRender.com. Figures 6A, 6E and 7A were created using PyMOL Molecular Graphics System, Version 3.0 Schrödinger, LLC.

## SUPPORTING INFORMATION

**S1 Figure: Expressing syt1 variants on WT background effect has minimal effect on evoked neurotransmission at 20Hz. (**A) Representative evoked inhibitory current traces of empty vector (CTRL) and expression of C2A syt1 variants. (B) Comparison of the mean eIPSC amplitude (nA) at 20Hz stimulation across C2A variants (Two-way ANOVA with Dunnett’s multiple comparisons test: CTRL vs. Syt1 OE p=0.994; CTRL vs. L159R p>999; CTRL vs. V184A p=772; CTRL vs. T196K p=0.960; CTRL vs. E209K p>0.999; CTRL vs. E219Q p>0.999; N=1-3, n=4-8 per group). (C) Comparison of paired-pulse ratio (PPR) at 20Hz stimulation for each group (One-way ANOVA with Dunnett’s multiple comparisons test: CTRL vs. Syt1 OE p>0.999; CTRL vs. L159R p=.765; CTRL vs. V184A p>0.999; CTRL vs. T196K p>0.999; CTRL vs. E209K p>0.999; CTRL vs. E219Q p=0.962; N=1-3, n=4-8 per group). (D) Representative evoked inhibitory current traces of empty vector (CTRL) and expression of C2B syt1 variants. (E) Comparison of mean eIPSC amplitude (nA) at 20Hz stimulation across C2B variants (Two-way ANOVA with Dunnett’s multiple comparisons test: CTRL vs. Syt1 OE p=0.994; CTRL vs. M303V p=.997; CTRL vs. D304G p=0.999; CTRL vs. S309P p=0.998; CTRL vs. N341S p=0.998; CTRL vs. Y365C p>0.999; CTRL vs. G369D p>0.999; N=1-3, n=4-8 per group). (F) Comparison of paired-pulse ratio (PPR) at 20Hz stimulation for each group (One-way ANOVA with Dunnett’s multiple comparisons test: CTRL vs. Syt1 OE p>0.999; CTRL vs. M303V p>0.999; CTRL vs. D304G p>0.999; CTRL vs. S309P p=0.487; CTRL vs. N341S p>0.999; CTRL vs. Y365C p=0.743; CTRL vs. G369D p>0.999; N=1-3, n=4-8 per group).

**S2 Figure: C2A and C2B variants effect on spontaneous excitatory neurotransmission on syt1KD background. (**A) Representative raw traces of baseline mEPSCs on Scramble (negative control), Syt1 knockdown (KD), Syt1 rescue (Syt1), and Syt1 variants in the C2A domain. (B) Comparison of baseline mEPSC amplitudes (pA) for each C2A variant (One-way ANOVA with Dunnett’s multiple comparisons test: SCR vs KD p=0.006; SCR vs Syt1 rescue p=0.369; SCR vs. L159R p=0.999; SCR vs. V184A p=0.477; SCR vs. T196K p=0.461; SCR vs. E209K p=0.995; SCR vs. E219Q p=0.794, N=2-5, n=6-19 per group). (C) Comparison of baseline mEPSC frequency (Hz) for each C2A variant (One-way ANOVA with Dunnett’s multiple comparisons test: SCR vs KD p<0.0001; SCR vs Syt1 rescue p=0.390; SCR vs. L159R p=0.987; SCR vs. V184A p=0.857; SCR vs. T196K p=0.533; SCR vs. E209K p=0.861; SCR vs. E219Q p>0.999, N=2-5, n=6-19 per group). (D) Representative raw traces of baseline mEPSCs on Scramble (negative control), Syt1 knockdown (KD), Syt1 rescue (Syt1), and Syt1 variants in the C2B domain. (E) Comparison of baseline mEPSC amplitudes (pA) on Scramble (negative control), Syt1 knockdown (KD), Syt1 rescue (Syt1), and Syt1 variants for each C2B variant (One-way ANOVA with Dunnett’s multiple comparisons test: SCR vs KD p=0.006; SCR vs Syt1 rescue p=0.369; SCR vs. M303V p=0.968; SCR vs. D304G p=0.044; SCR vs. S309P p=0.038; SCR vs. N341S p<0.0001; SCR vs. Y365C p=0.278; SCR vs. G369D p=0.706, N=2-5, n=6-19 per group). (F) Comparison of baseline mEPSC on Scramble (negative control), Syt1 knockdown (KD), Syt1 rescue (Syt1), and Syt1 variants in the C2B domain frequency (Hz) (One-way ANOVA with Dunnett’s multiple comparisons test: SCR vs KD p<0.0001; SCR vs Syt1 rescue p=0.390; SCR vs. M303V p>0.999; SCR vs. D304G p=0.610; SCR vs. S309P p=0.998; SCR vs. N341S p<0.0001; SCR vs. Y365C p=0.969; SCR vs. G369D p>0.999, N=2-5, n=6-19 per group).

**S3 Figure: C2A and C2B variants effect on spontaneous inhibitory neurotransmission on Syt1 KD background.** (A) Representative raw traces of baseline mIPSCs on Scramble (negative control), Syt1 knockdown (KD), Syt1 rescue (Syt1), and Syt1 variants in the C2A domain. (B) Comparison of baseline mIPSC amplitudes (pA) for each C2A variant (One-way ANOVA with Dunnett’s multiple comparisons test: SCR vs KD p=0.999; SCR vs Syt1 rescue p=0.636; SCR vs. L159R p=0.999; SCR vs. V184A p=0.489; SCR vs. T196K p=0.024; SCR vs. E209K p>0.999; SCR vs. E219Q p=0.362, N=1-3, n=3-10 per group). (C) Comparison of baseline mIPSC frequency (Hz) for each C2A variant (One-way ANOVA with Dunnett’s multiple comparisons test: SCR vs KD p=0.009; SCR vs Syt1 rescue p=0.587; SCR vs. L159R p=0.991; SCR vs. V184A p=0.018; SCR vs. T196K p<0.0001; SCR vs. E209K p=.997; SCR vs. E219Q p=0.786, N=1-3, n=3-10 per group). (D) Representative raw traces of baseline mIPSCs on Scramble (negative control), Syt1 knockdown (KD), Syt1 rescue (Syt1), and Syt1 variants in the C2B domain. (E) Comparison of baseline mIPSC amplitudes (pA) on Scramble (negative control), Syt1 knockdown (KD), Syt1 rescue (Syt1), and Syt1 variants for each C2B variant (One-way ANOVA with Dunn’s multiple comparisons test: SCR vs KD p>0.999; SCR vs Syt1 rescue p>0.999; SCR vs. M303V p=0.110; SCR vs. D304G p>0.999; SCR vs. S309P p>0.999; SCR vs. N341S p=0.027; SCR vs. Y365C p=0.027; SCR vs. G369D p>0.999, N=1-3, n=3-10 per group). (F) Comparison of baseline mIPSC frequency (Hz) for each group on Scramble (negative control), Syt1 knockdown (KD), Syt1 rescue (Syt1), and Syt1 variants in the C2B domain (One-way ANOVA with Dunn’s multiple comparisons test: SCR vs KD p=0.035; SCR vs Syt1 rescue p=0.683; SCR vs. M303V p>0.999; SCR vs. D304G p=0.400; SCR vs. S309P p>0.999; SCR vs. N341S p<0.0001; SCR vs. Y365C p=0.019; SCR vs. G369D p>0.999, N=1-3, n=3-10 per group).

**S4 Figure: Selective MEK I/II inhibitors do not impact spontaneous release frequency in N341S variant networks. (**A) Representative raw traces of mEPSCs of empty Vector control (CTRL) and syt1-mutated disease variant (N341S) disease variant baseline and after acute 1hr treatment with either 1µM Trametinib or 10µM U0126. (B) Comparison of mean baseline mEPSC amplitude (pA) for each group (Two-way ANOVA with Sidak’s multiple comparisons test [DMSO versus Trametinib]: [CTRL] DMSO vs. Trametinib p=0.968; [N341S] DMSO vs. Trametinib p>0.999; [CTRL] DMSO vs. U0126 p=0.941; [N341S] DMSO vs. U0126 p=0.886, N=2, n=6-8 per group). (C) Comparison of mean baseline mEPSC frequency (Hz) for each group (Two-way ANOVA with Sidak’s multiple comparisons test [DMSO versus Trametinib]: [CTRL] DMSO vs. Trametinib p=0.999; [N341S] DMSO vs. Trametinib p=0.892; [CTRL] DMSO vs. U0126 p=0.999; [N341S] DMSO vs. U0126 p=0.600, N=2, n=6-8 per group).

